# Allosteric activation of CRISPR-Cas12a requires the concerted movement of the bridge helix and helix 1 of the RuvC II domain

**DOI:** 10.1101/2022.03.15.484427

**Authors:** Elisabeth Wörle, Anthony Newman, Gaetan Burgio, Dina Grohmann

**Author notes:** For correspondence: Dina Grohmann, Department of Biochemistry, Genetics and Microbiology, Institute of Microbiology, University of Regensburg, Universitätsstraße 31, 93053 Regensburg, Germany, E-Mail.

## Abstract

Nucleases derived from the prokaryotic defense system CRISPR-Cas are frequently re-purposed for gene editing and molecular diagnostics. Hence, an in-depth understanding of the molecular mechanisms of these enzymes is of crucial importance. We focused on Cas12a from *Francisella novicida* (FnCas12a) and investigated the functional role of helix 1, a structural element that together with the bridge helix (BH) connects the recognition and the nuclease lobes of FnCas12a. Helix 1 is structurally connected to the lid domain that opens upon DNA target loading thereby activating the active site of FnCas12a. We probed the structural states of FnCas12a variants altered in helix 1 and/or the BH using single-molecule FRET measurements and assayed the pre-crRNA processing, *cis-* and *trans-*DNA cleavage activity. We show that helix 1 and not the BH is the predominant structural element that confers conformational stability of FnCas12a. Even small perturbations in helix 1 lead to a decrease in DNA cleavage activity while the structural integrity is not affected. Our data, therefore, implicate that the concerted remodeling of helix 1 and the BH upon DNA binding is structurally linked to the opening of the lid and therefore involved in the allosteric activation of the active site.

## Introduction

CRISPR-Cas (CRISPR – <u>c</u>lustered regularly interspaced <u>s</u>hort <u>p</u>alindromic repeats, Cas – <u>C</u>RISPR <u>as</u>sociated) endonucleases are crucially involved in the adaptive anti-phage immune system in archaea and bacteria (1–6). This includes a wide range of DNA and/or RNA nucleases that are involved in spacer recruitment, processing, and integration into the CRISPR locus in the genome (Cas1/2 and Cas4), over RNA nucleases that process the primary transcript, the CRISPR RNA (crRNA), to effector nucleases that eventually cleave target DNA or RNA (7–10). Single effector nucleases like Cas9 or Cas12a have gained special attention as they can be easily programmed with a crRNA to recognize and cleave specific target nucleic acid sequences (11, 12). Based on the protein repertoire encoded by a CRISPR-Cas locus and the chemistry of the target nucleic acid (DNA or RNA), CRISPR-Cas systems are classified into two classes and further divided into six types and a constantly growing number of subtypes (2, 7, 8). The single-effector nuclease Cas12a, subject of this study, is a class 2 protein of the type V-A family (12, 13).

To avoid self-targeting and to exclusively target foreign sequences, Cas12a recognizes a protospacer adjacent motif (PAM, 5’-TTTV-3’) in the target DNA (14–16). After binding, R-loop formation occurs with the crRNA forming an RNA-DNA hybrid with the complementary target strand (TS) of the target DNA (15, 17, 18). The non-target strand (NTS) is directly guided to the RuvC domain harboring the single Cas12a nuclease active site. Consequently, the target DNA is sequentially cleaved with the NTS being cleaved 20 times faster than the TS (18, 19). Ultimately, a double-strand-break (DSB) with staggered ends (5-8 nt 5’ overhang) is produced by Cas12a (12, 18–20). After cleavage, the PAM-distal product is released whereas Cas12a remains bound to the PAM-proximal fragment (21–23) and further trims the NTS in a stepwise manner (19, 24). The dissociation of the cleaved PAM-proximal part is very slow and consequently, Cas12a is a single-turnover enzyme (22).

crRNA-directed cleavage of target DNA (called *cis*-cleavage) activates Cas12a for non-specific cleavage of ssDNA, ssRNA, and dsDNA (25–29). This so-called *trans*-cleavage activity is not completely understood (30) but has already been exploited to design quantitative detection platforms for nucleic acids (31, 32).

The bilobal structure of Cas12a can be divided into the nuclease (Nuc) lobe (composed of the RuvC I, RuvC II, RuvC III, BH (bridge helix), Nuc domains), and the recognition (REC) lobe (composed of the REC1 and REC2 domains). Both lobes are connected by the wedge domain (WED I, WED II, WED III, PI (<u>P</u>AM interacting)) that acts as a hinge (11, 15, 33, 34). The wedge domain also anchors the pseudoknot structure that is formed by the 5’-end of the crRNA. The crRNA furthermore interacts with the RuvC and REC domains (11, 23, 33, 34). Upon loading of target DNA to the Cas12a-crRNA complex, the Nuc and REC lobes transition from a closed to an open conformation (11, 15, 24).

The BH is a structural element that links the Nuc and the REC lobe and interacts with the sugar-phosphate backbone of the TS. Furthermore, structural studies of *Acidaminococcus sp*. Cas12a (AsCas12a) showed that the BH is involved in the recognition of the crRNA-DNA-hybrid (34) whereas only a few contacts between the BH residues lysine 978, isoleucine 977, and asparagine 976 of FnCas12a and the template strand were described (23). The BH harbors a tryptophan residue (W971 in Cas12a encoded by *F. novicida*) that is buried in a hydrophobic pocket of the REC2 domain and thereby stabilizes the active conformation of Cas12a (23, 34). Previous studies showed that mutations in the BH have drastic effects on the cleavage activity and accuracy of Cas12a (24, 35). However, FnCas12a variants with full deletion of the BH are still able to bind crRNA and target DNA and to undergo the conformational transition upon crRNA and DNA binding (24). According to crystal structures of the binary Cas12a-crRNA complex and the ternary Cas12a-crRNA-target DNA complex (11, 15, 18, 23, 33, 34, 36), the BH is connected to and acts in concert with the adjacent helix of the RuvC domain (termed helix 1). In the binary complex the BH is short, whereas helix 1 is long: In contrast, the BH is elongated in the ternary complex but helix 1 is shortened by six amino acids (**Figure 1**). This tandem movement ensures that the tryptophan residue (W971) can stay in its hydrophobic binding pocket in the REC domain that undergoes a re-positioning upon the transition from binary to ternary complex.

**Figure 1.**
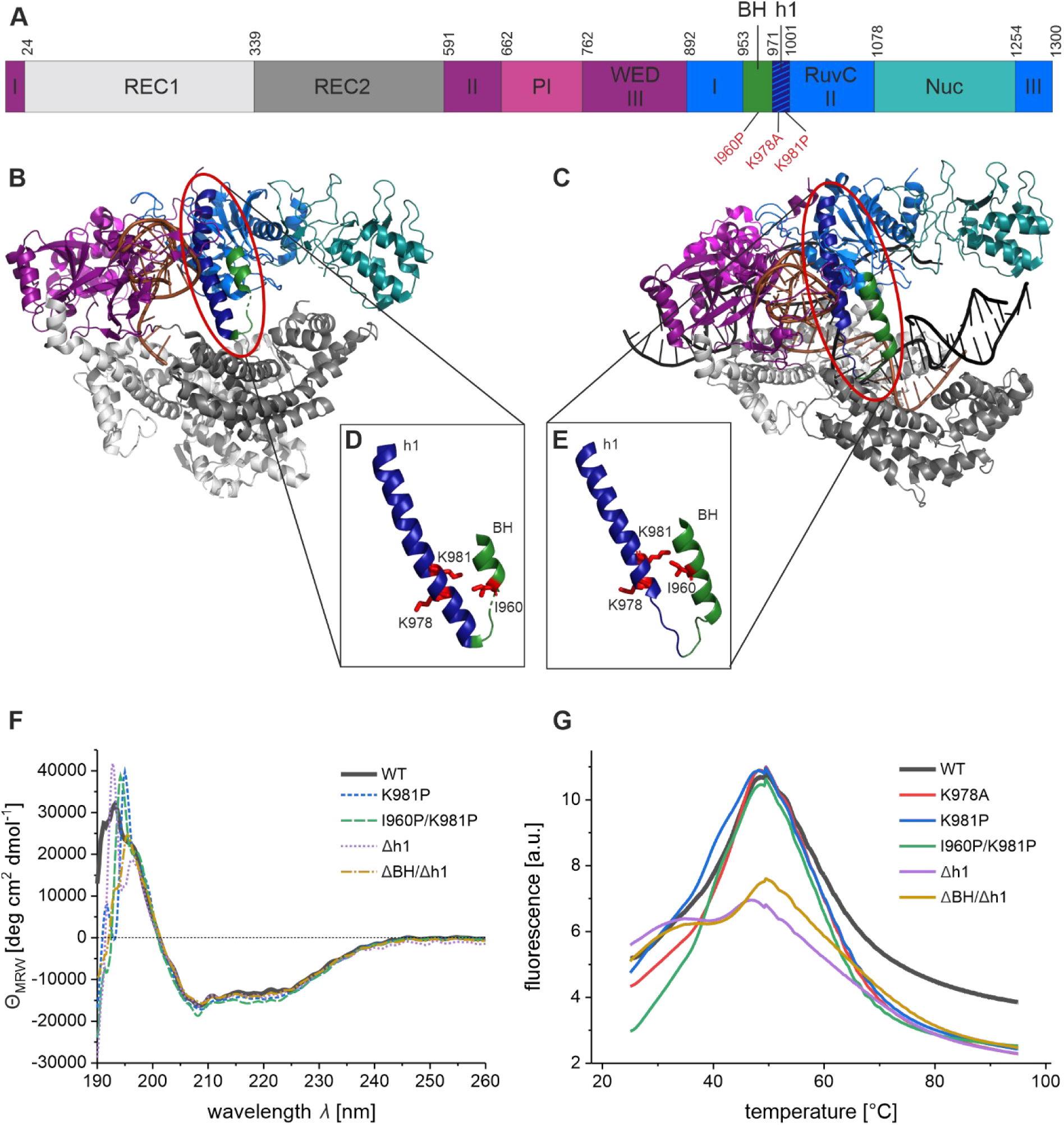
Structural organization of *Francisella novicida* Cas12a (FnCas12a) and influence of helix 1 mutations on the overall structure and thermo stability. (**A**) Domain organization of FnCas12a with mutation sites indicated in red. (**B**) Structure of the binary complex of FnCas12a with crRNA (orange) (PDB: 5NG6). (**C**) Structure of the ternary complex of FnCas12a together with crRNA (orange) and target DNA (black) (PDB: 6I1K). (**B**) and (**C**) Protein domains are colored according to the domain organization in (A). Helix 1, a part of RuvC II domain, is colored in dark blue, the bridge helix (BH) in green. (**D**) and (**E**) Close-up of helix 1 (blue) and BH (green) in the binary (D) and ternary (E) state, showing the change of helical lengths upon the rearrangement from binary to ternary complex. Mutation sites I960, K978, and K981 are shown in red. (**F**) Far-UV CD spectra of WT FnCas12a and FnCas12a helix 1 variants (2 µM) at 25 °C. Shown is the average of five replicates. (**G**) Protein Thermal Shift™ (Thermo Scientific) melting curves of FnCas12a variants (2 µg) from 25 to 95 °C. Shown is the average of four replicates.

In this study, we generated FnCas12a variants with distorted (K981P and I960P/K981P) or deleted helix 1 (Δh1 and ΔBH/Δh1) and sought to understand whether the presence and integrity of helix 1 are required for (i) crRNA binding and pre-crRNA processing, (ii) DNA binding and *cis*-cleavage, (iii) *trans*-cleavage and (iv) conformational changes of FnCas12a during its activity cycle.

Our study reveals that deletion of helix 1 abrogates the *cis-* and *trans*-DNA cleavage activity of FnCas12a while the pre-crRNA processing activity remains unaffected. Single-molecule FRET analysis demonstrates that the closure of FnCas12a during the binary to ternary complex transition cannot commence in the absence of helix 1, explaining the severe impact on the DNA cleavage activity. Notably, even small disturbances in the helix 1-BH tandem movement severely impact the DNA cleavage activities suggesting that the fine-tuning of the helical structure of helix 1 and BH have a direct impact on the opening and closure of the adjacent lid domain and by extension on the DNA cleavage activity as an opening of the lid is a prerequisite for the activation of the catalytic site in the RuvC domain. Taken together, our data show that in FnCas12a – unlike in Cas9 – not the BH but helix 1 is the main structural element. Helix 1 not only connects the Rec and Nuc lobe but influences the allosteric activation of the RuvC domain via the helix 1-BH tandem motion at the ternary complex formation step.

## Material and Methods

### Protein expression and purification

Wildtype FnCas12a and FnCas12a variants were expressed and purified as previously described (24). In short, the Cas12a gene from *Francisella novicida* (Addgene #69975) was cloned into a pGEX-2TK expression vector. FnCas12a variants were generated via site-directed mutagenesis (for mutagenesis primers see Supplementary **Table S1**). The proteins were expressed in *E. coli* BL21(DE3) cells. After cell lysis and clearance of the lysate, proteins were purified via an N-terminal GST-tag. The GST-tag was removed utilizing the thrombin protease cleavage site (100 U/ml, 50:1 (v/v)) and overnight incubation at 4 °C). FnCas12a was further purified via heparin affinity chromatography. Protein concentrations were calculated using an extinction coefficient at 280 nm of 143 830 M^−1^cm^−1^ (ProtParam tool from www.expasy.org). The protein storage buffer was composed of (20 mM Tris-HCl pH 7.5, 50 mM NaCl, 5 mM MgCl_2_, 5% (v/v) glycerol) and protein aliquots were flash-frozen and stored at −80 °C until further use.

### Site-specific incorporation of the unnatural amino acid *para*-azido-L-phenylalanine into FnCas12a

In order to prepare site-specifically labeled FnCas12a for single-molecule measurement, Cas12a variants with site-specifically incorporated unnatural amino acid *para*-azido-L-phenylalanine (AzF) were generated via the Amber-suppressor strategy (37, 38) as previously described (24). AzF was incorporated at positions D470 and T1222 (for mutagenesis primers see Supplementary **Table S1**). Plasmids that encoded the Amber codons at the chosen site of the FnCas12a gene were expressed in BL21(DE3)-pEVOL-pAzF, cells with an additional plasmid encoding for an arabinose-inducible promoter to express Amber-suppressor tRNA (tRNA^CUA^) and a biorthogonal tRNA synthetase (Addgene #31186)(37). During expression, *E. coli* cells were supplemented with 200 mg/ml of AzF (ChemImpex). AzF-modified proteins were purified as described above.

### Protein labeling

As described previously (24), AzF-modified FnCas12a variants were stochastically labeled with DyLight550 and DyLight650 via the Staudinger-Bertozzi ligation (39). The labeling reaction of the purified protein was conducted with a final concentration of 50 µM of DyLight550 and DyLight650 for 90 min at room temperature in darkness. The labeled protein was purified via Sephadex G-50 illustra NICK size exclusion chromatography column using Cas batch buffer (50 mM Tris-HCl pH 7.5, 200 mM NaCl, 5 mM MgCl_2_, 2% (v/v) glycerol, 0.1% (v/v) Tween20) and was directly used for single-molecule measurements.

### Synthetic oligonucleotides

The target DNA sequence was derived from Swarts *et al*. (TS 5’-ACTCAATTTTGACAGCCCACATGGCATTCCACTTATCACTAAAGGCATCCTTCCACGT-3’, NTS 5’-ACGTGGAAGGATGCCTTTAGTGATAAGTGGAATGCCATGTGGGCTGTCAAAATTGAGT-3’) (40). The matched target MM 0 as well as the mismatched target DNA sequences (MM 2, MM 10, MM 20), were purchased from Eurofins-MWG (Ebersberg, Germany, **Table S2**). For cleavage assays, the target strand carried a Cy5 label, the non-target strand a Cy3 label either at the 5’- or the 3’-end of the DNA. Annealing was carried out as described in Wörle *et al*. (24).

Cy5-labeled crRNA (5’-AAUUUCUACUGUUGUAGAUGUGAUAAGUGGAAUGCCAUGUGGG-Cy5-3’) was purchased from Microsynth Seqlab GmbH (Göttingen, Germany) and Atto532 labeled pre-crRNA (5’-Atto532-UAAUUUCUACUGUUGUAGAUGUGAUAAGUGGAAUGCCAUGUGGG-3’) was purchased from Biomers (Ulm, Germany).

### *In vitro* transcription

The crRNA (5’-AAUUUCUACUGUUGUAGAUGUGAUAAGUGGAAUGCCAUGUGGG-3’) was synthesized and purified using the T7 RiboMAX™ Express Large Scale RNA Production System (Promega). The following DNA templates were used for the *in vitro* transcription reaction: T7crRNA 5’-CCCACATGGCATTCCACTTATCACATCTACAACAGTAGAAATTCCCTATAGTGAGTCGTATTATCGATC-3’ and T7 promotor oligo 5’-GATCGATAATACGACTCACTATAGGG-3’ and were purchased from Eurofins-MWG (Ebersberg, Germany).

### Electrophoretic mobility shift assay

crRNA-Cas12a complexes for electrophoretic mobility shift assays were formed in 1x Cas buffer (20 mM Tris-HCl pH 7.5, 100 mM NaCl, 10 mM MgCl_2_, 2% (v/v) glycerol, 1 mM DTT, 0.05% (v/v) Tween20) with a two-fold excess of protein (20 nM) over fluorescently labeled crRNA (10 nM). The complexes were incubated in a total volume of 10 µl at room temperature for 10 min. Gel loading buffer (final concentration: 62.5 mM Tris-HCl pH 6.8, 2.5% (w/v) Ficoll 400) was added and the samples separated on a non-denaturing TBE gel (15% PAA, 300 V, 10 min). In-gel fluorescence of the labeled crRNA was visualized using a ChemiDoc Imaging System (Bio-Rad).

Electrophoretic mobility shift assays of ternary complexes (Cas12a-crRNA-target DNA) were conducted as previously described (24). The pre-formed binary complex (200 nM) was added in 7.5-fold (75 nM) excess to 10 nM of fluorescently labeled target DNA (10 nM). After 1 h reaction time at 37 °C, 32 g/ml Heparin was added to avoid unspecific binding. After 10 min incubation at room temperature, gel loading dye (final concentration: 62.5 mM Tris-HCl pH 6.8, 2.5% (w/v) Ficoll 400) was added and the samples separated by non-denaturing Tris-HCl gel electrophoresis (6% PAA, 230 V, 30 min). The in-gel fluorescence was visualized using a ChemiDoc Imaging System (Bio-Rad).

### Plasmid cleavage assay

Plasmid cleavage assays were performed using the eGFP-hAgo2 plasmid (Addgene #21981) as target DNA, either as supercoiled or linearized plasmid (linearization with SmaI). Cleavage assays were carried out as previously described (24). In brief, the binary complex composed of FnCas12a (100 nM) and crRNA (100 nM) was pre-formed and incubated at room temperature for 10 min. 37.5 nM of binary complexes were added to 5 nM of target DNA and incubated for 1 h at 37 °C. To stop the reaction, 0.36 U Proteinase K (Thermo Fisher) was added and incubated for 30 min at 55 °C. The reaction with the linearized plasmid was analyzed on a 0.8% agarose 1× TAE gel, the reaction of the supercoiled DNA on a 1.5% agarose 1x TAE gel and visualized using SYBR-Safe (Invitrogen).

### DNA cleavage assay using a short DNA target

Cleavage assays were conducted as previously described (24) using 7.5-fold excess of protein and crRNA (final concentration of 75 nM) over fluorescently labeled target DNA (10 nM). The reactions were incubated for 1 h at 37 °C. To stop the cleavage reaction, 0.36 U Proteinase K (Thermo Fisher) was added, and the reaction was incubated at 55 °C for 30 min. Samples were mixed with loading dye (final concentration: 47.5% (v/v) formamide, 0.01% (w/v) SDS, 0.01% (w/v) bromophenol blue, 0.005% (w/v) xylene cyanol, 0.5 mM EDTA), heated to 95 °C and loaded onto a pre-heated 15% PAA, 6 M Urea, TBE gel (300 V, 30min). Fluorescent signals were visualized using a ChemiDoc Imaging System (Bio-Rad).

Cleavage reactions that were resolved on high-resolution sequencing gels (15% PAA, 6 M Urea, 1× TBE) contained 750 nM of pre-formed binary complex and 100 nM of fluorescently labeled target DNA. The samples were separated on a pre-heated 45 cm sequencing gel (45 °C, 65 W, 1 h) and visualized using a ChemiDoc Imaging System (Bio-Rad).

### *Trans*-cleavage using single-stranded DNA

First, FnCas12a ribonucleoprotein complexes were formed that prime FnCas12a for *cis*-cleavage. 1 µM FnCas12a and 1 µM crRNA were complexed at 25 °C for 10 min in DEPC-treated H_2_O. Afterwards, 50 nM of a single-stranded DNA oligo ‘activator’ were added and the sample was incubated at 30 °C for 20 min in *trans*-cleavage buffer (10 mM Tris-HCl pH 7.9, 100 mM NaCl, 10 mM MgCl_2_, 50 µg/ml BSA). To detect *trans*-cleavage activity, the *cis*-cleavage reaction was diluted 10x in buffer to a total volume of 150 µl, three technical replicates of 50 µl for each condition were loaded onto a 96-well plate (black with clear bottom, Invitrogen). 50 µl of 200 nM fluorescence quencher (FQ) substrate (5’FAM-TTTTTTTTTT-3’IABk, IDT, diluted in 1xbuffer) were added to each well, the plate was immediately transferred to a plate reader (Victor Nivo Multimode, PerkinElmer), shaken at 300 rpm (orbital) for 30 s, and fluorescence intensity measured over 90 min (excitation filter 480/30 nm, emission filter 530/30 nm). The final concentrations were as follows: 100 nM Cas12a RNP, 5 nM ssDNA, and 100 nM FQ reporter, and upper limit of 5 nM enzyme. Using WT FnCas12a and variants at variable concentrations of FQ reporter, a calibration curve was constructed to convert changes in fluorescence intensity to *trans-*cleavage rate and concentration of cleaved FQ reporter over time. The slope of change in fluorescence intensity was extracted from the first 600 seconds of the reaction, and relative rates of *trans-*cleavage estimated from a calibration curve. Michaelis-Menten Kinetics (v_m_, K_M_, k_cat_ and K_M_/k_cat_) were calculated using a similar approach described in Ramachandran et al. 2021 with modification (41). Briefly, the background-subtracted maximum fluorescence intensity was retained to calculate the maximal velocity and the concentration of cleaved FQ over time. Velocity was determined as the ratio between the total fluorescence intensity (F(t)), the initial FQ concentration (c_0_) the fluorescence levels of uncleaved reporter (F_ucl_ - FQ without RNP complex) and the cleaved reporter (F_cl_ - FQ in presence of the RNP complex and its cognate DNA target) as followed:

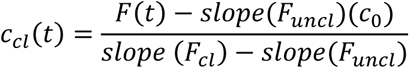

Kinetics parameters were obtained by fitting the calibration curves and k_cat_ and K_M_ were calculated using GraphPad Prism Software (GraphPad, CA, USA).

### Pre-crRNA processing assay

Pre-crRNA processing was shown with an electrophoretic mobility shift assay of the pre-crRNA with FnCas12a. The protein-pre-crRNA complexes were formed in 1x Cas buffer (20 mM Tris-HCl pH 7.5, 100 mM NaCl, 10 mM MgCl_2_, 2% (v/v) glycerol, 1 mM DTT, 0.05% (v/v) Tween20) with a two-fold excess of protein (20 nM) over 5’ fluorescently labeled pre-crRNA (10 nM). The complexes were incubated in a total volume of 10 µl at room temperature for 10 min. Gel loading buffer (final concentration: 62.5 mM Tris-HCl pH 6.8, 2.5% (w/v) Ficoll 400) was added and the samples separated on a non-denaturing TBE gel (15% PAA, 300 V, 10 min). In-gel fluorescence of the labeled crRNA was visualized using a ChemiDoc Imaging System (Bio-Rad).

### Protein Thermal Shift™ melting curves

Melting curves of FnCas12a variants were obtained using the Protein Thermal Shift™ kit (ThermoFisher) using the Rotor-Gene Q qPCR cycler (Qiagen) and the HRM (High-Resolution Melt) software. The proteins were diluted in water and measurements were performed in quadruplicates. The data were analyzed using the Origin software (ADDITIVE Friedrichsdorf, Germany).

### Circular dichroism spectroscopy (CD)

CD spectra were recorded at a wavelength range from 190 to 260 nm. Samples were measured in 1x Cas buffer (20 mM Tris-HCl pH 7.5, 100 mM NaCl, 10 mM MgCl_2_, 2% (v/v) glycerol, 1 mM DTT, 0.05% (v/v) Tween20) in a quartz crystal cuvette (0.2 mm, HELLMA) using the spectropolarimeter J-815 (JASCO). The molecular ellipticity per amino acid was obtained by normalization according to the following equation:

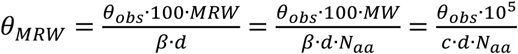

With *ϑ*_*MRW*_ as the average ellipticity per amino acid (deg cm^2^ dmol^−1^), *ϑ*_*obs*_ the measured ellipticity (mdeg), *MRW* the mean residue weight (in kDa), *β* the protein concentration in mg/ml, *c* the protein concentration in µmol/l, *d* the thickness of the cuvette (cm), *MW* the molecular weight of the protein (in kDa), and *N*_*aa*_ the number of amino acids of the protein.

### Confocal single-molecule FRET measurements and data analysis

Single-molecule FRET measurements in solution were conducted as previously described (24) using a MicroTime 200 confocal microscope (PicoQuant) with pulsed laser diodes (532 nm: LDH-P-FA-530B; 636 nm: LDH-D-C-640; PicoQuant / clean-up filter: ZET 635; Chroma). The data were analyzed using the software package PAM (42) and fitted with two or three Gaussian equations using the Origin software (ADDITIVE Friedrichsdorf, Germany).

### Structure prediction with Alphafold2

Structural prediction was performed using the non-docker version of Alphafold2 (43) available at (https://github.com/kalininalab/alphafold_non_docker) with the default settings and no custom multiple sequence alignment (MSA) template as a default option. The input sequences were obtained from FnCas12a crystal structure bound to the crRNA and target DNA (Protein Data Bank - PDB 6I1K) and then modified with the respective mutations. The accuracy of the structural model was tested using a per-residue measure of local confidence (predicted Local Distance Difference Test, pLDDT) and indicated a high confidence in accurately predicting the residues and domains. Structures were analyzed with PyMol version 2.3.1 (Schrodinger LLC).

## Results

### Structural characterization of FnCas12a helix 1 mutations and deletion variants

Recently, we characterized Cas12a variants from *Francisella novicida* that carried mutations in or a full deletion of the bridge helix (BH) element. This study revealed that the overall structural integrity of the protein was conserved even in the case of the BH deletion variant, but mutations affect the trimming activity, mismatch sensitivity, and cleavage rate of FnCas12a (24). While the BH constitutes the main structural element that connects the Nuc and REC lobe in Cas9 proteins (44–48), the BH is connected to an adjacent helix in Cas12a proteins (termed helix 1) (11, 15, 24, 33, 34, 36). Here, we investigated to which extent helix 1 contributes to the stability and activity of FnCas12a. We, therefore, created a series of FnCas12a variants that carry mutations and/or deletions in helix 1 only or double mutations in the BH and helix 1 (**Figure 1 A-C**). This includes point mutations in helix 1 (K981P) or double mutations in the BH and helix 1 (I960P/K981P) designed to break the alpha-helical structure of the BH and helix 1. Moreover, we introduced a single mutation at position K978 in helix 1 (K978A) to interrupt an interaction of the positively charged lysine side chain with the backbone of the template DNA. We furthermore created a helix 1 deletion variant (Δh1, deletion of residues K972-N1000) and a variant in which the BH and helix 1 are deleted (ΔBH/Δh1, deletion of residues Y953-N1000). Structural predictions using the AlphaFold2 algorithm showed no significant structural changes in FnCas12a when these mutations are introduced (**Figure S1, Table S3**). For the K978A, K981P, and I960P/K981P variants, homogeneous protein preparations were obtained. However, following the same expression and purification protocol, the Δh1 and ΔBH/Δh1 preparations were less homogeneous indicating that the protein variants might be less stable than the wildtype and single mutants (**Figure S2**). In a series of biophysical experiments, we tested whether the mutations and/or deletions influence the overall stability or structural organization of FnCas12a. Circular dichroism experiments revealed that all variants exhibit a minimum at 208 nm and a less pronounced minimum at 222 nm as well as a maximum at 192-198 nm in the far-UV spectrum indicative of proteins with a predominantly alpha-helical structure (**Figure 1F**). Indeed, FnCas12a is almost exclusively composed of alpha helices and the FnCas12a variants appear not to vary from this overall structural organization. However, protein thermal shift assays revealed an additional thermal melting point at 35.9 °C for the Δh1 and ΔBH/Δh1 variants (**Figure 1G**). For WT FnCas12a and the K978A, K981P, I960P/K981P, and the ΔBH variants only a single melting point at 48.2 °C was observed (**Figure 1G** and **S3**). These data indicate that the deletion of helix 1 leads to a partial destabilization of the protein.

Previously, we elucidated the conformational transition of FnCas12a during the activity cycle using smFRET measurements (24). In brief: the apo form of wildtype FnCas12a adopts an open (low FRET) and closed (high FRET) state. Addition of the crRNA and formation of the binary crRNA-Cas12a complex shifts this equilibrium completely towards the closed state. The transition from the binary to the ternary complex is accompanied by a structural rearrangement from a closed (mean FRET efficiency of E=0.97) to a more open state (mean FRET efficiency of E=0.82) of the enzyme (**Figure 2**) (24). In order to follow the conformational states and transitions of the FnCas12a variants used in this study, incorporated unnatural amino acids for the site-specific labeling of the variants with labeling positions at D470 and T1222 in the REC and Nuc lobe, respectively (**Figure S4**). As reported previously, the apo form of WT FnCas12a adopts an open (population with a mean FRET efficiency of E=0.12) and closed (population with a mean FRET efficiency of E=0.97). Similarly, the I960P/K981P variant adopts the open and closed state. However, the Δh1 and ΔBH/Δh1 resulted in an almost complete loss of the high FRET population indicating that these FnCas12a variants cannot adopt the closed state (**Figure 2**). Furthermore, the addition of crRNA did not induce the conformational change towards the closed conformation in the case of the Δh1 and ΔBH/Δh1 variants (**Figure 2**) neither did the addition of crRNA and target DNA induce any changes in the FRET distribution. This is most likely a result of the defect in crRNA and dsDNA binding of these FnCas12a variants that lack helix 1. Compared to the wildtype enzyme, no significant differences in the FRET efficiency histograms for the apo enzyme, binary, or ternary complex were observed for the FnCas12a^I960P/K981P^ variant indicating that this variant efficiently undergoes the open to closed (apo → binary complex) and closed to semi-closed (binary → ternary complex) transitions.

**Figure 2.**
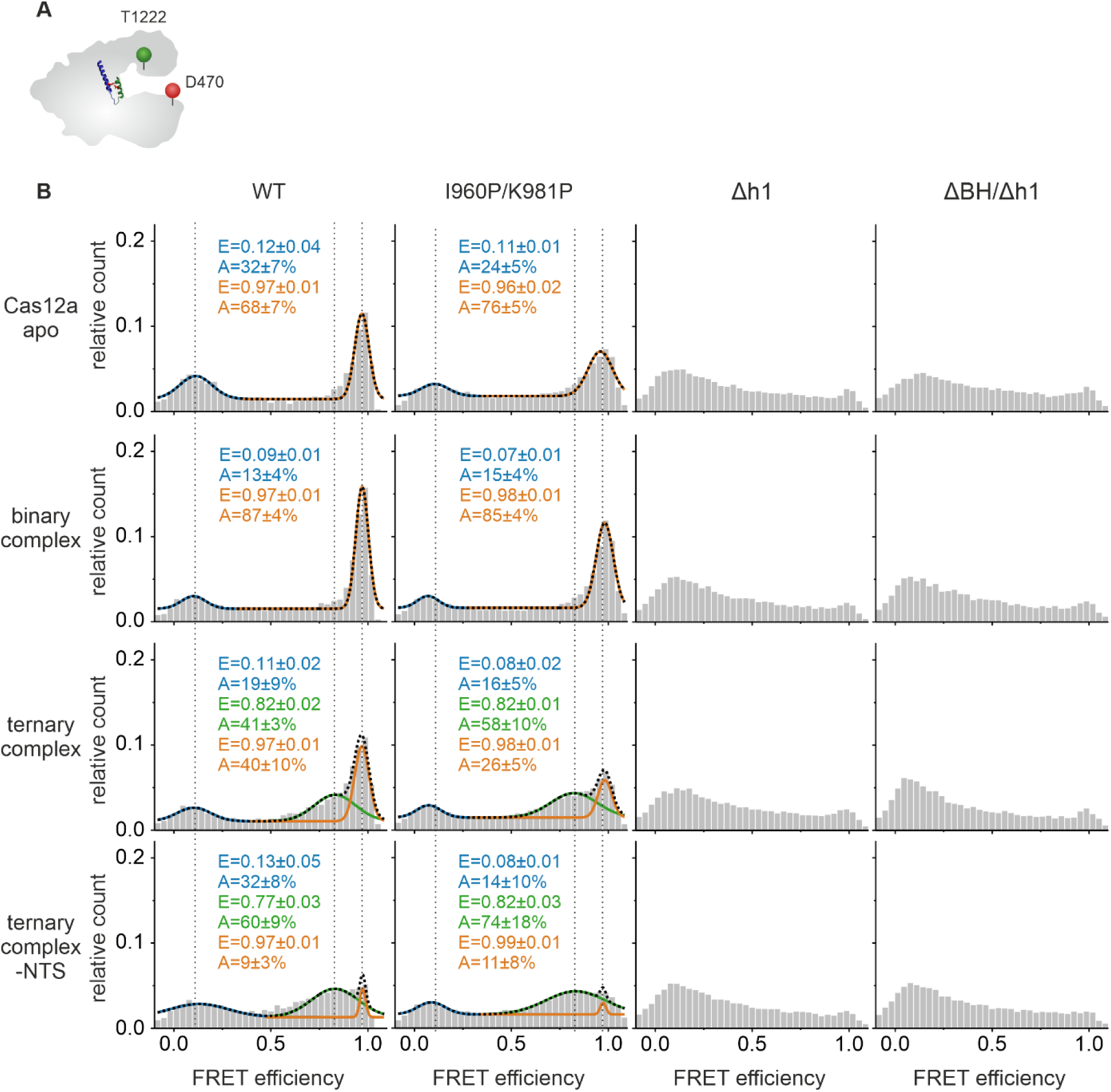
Single molecule Förster resonance energy transfer (FRET) measurements of freely diffusing molecules were conducted of doubly labeled FnCas12a^REC-Nuc*DL550/DL650^ helix 1 variants in comparison to WT FnCas12a^REC-Nuc*DL550/DL650^. (**A**) Schematic illustration of FnCas12a with labeling positions in the Nuc domain (T1222) and the REC domain (D470). (**B**) FRET efficiency histograms of FnCas12a I960P/K981P^REC-Nuc*DL550/DL650^, FnCas12a Δh1^REC-Nuc*DL550/DL650^, and FnCas12a ΔBH/Δh1REC-Nuc*DL550/DL650 compared to WT FnCas12a^REC-Nuc*DL550/DL650^ of the apo enzyme, the binary complex (1 nM crRNA), the ternary complex (1 nM crRNA, 1 nM target DNA), and the ternary complex without NTS (1 nM crRNA, 1 nM TS DNA). Fitting of the histograms was performed with a double or triple Gaussian function. The averaged mean FRET efficiencies (E) and the percentage distribution of the populations (A) are given with SEs in the histograms. Measurements were performed in triplicates (see also Supplementary Figure S8).

These findings indicate that loss of helix 1 results in a conformationally destabilized FnCas12a. Hence, we conclude that helix 1 is crucially involved in maintaining the structural integrity of FnCas12a.

### FnCas12a-helix 1 variants bind and process pre-crRNAs

As crRNA addition did not induce the conformational transition of FnCas12a helix 1 variants to the closed conformation, we next tested their ability to bind and process crRNA. Electrophoretic mobility shift assays using fluorescently labeled crRNA demonstrate that FnCas12a variants that carry a single mutation in helix 1 efficiently bind the crRNA (**Figure 3A/B**) albeit at slightly reduced levels as compared to the wildtype (**Figure 3C**). An exception is the K978A variant that binds crRNAs as efficient as WT FnCas12a. The Δh1 and ΔBH/Δh1 variants are impaired in their capacity to bind crRNA (Δh1 10.7%, ΔBH/Δh1: 22.9% binding efficiency relative to wildtype FnCas12a). Notably, the ΔBH variant does still bind the crRNA albeit with 50% reduced efficiency as compared to the wildtype indicating that deletion of helix 1 leads to a drastic change in crRNA binding capacity.

**Figure 3.**
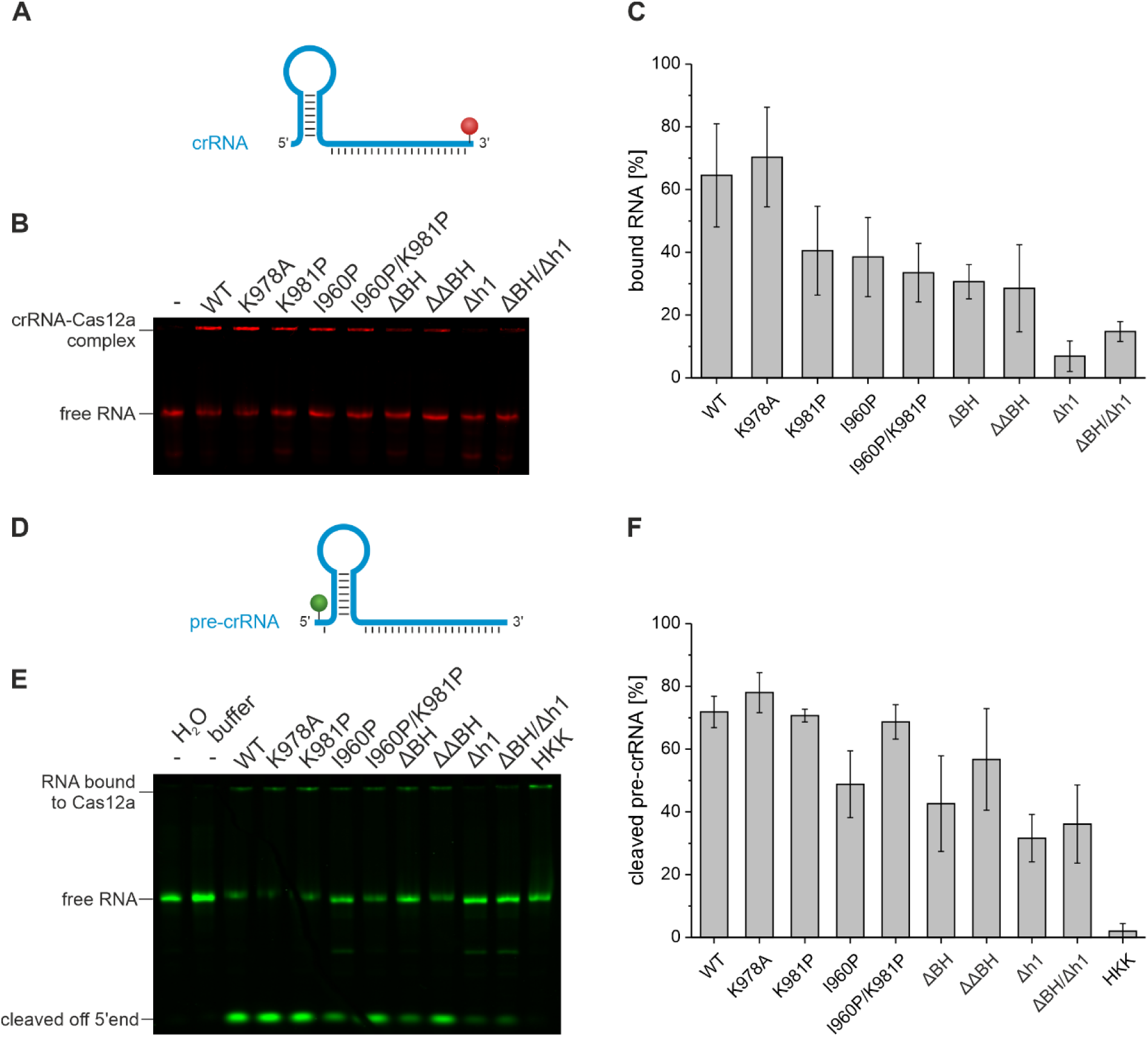
Binding behavior to crRNA and pre-crRNA processing of FnCas12a helix 1 variants in comparison to FnCas12a BH variants. (**A**) Schematic representation of crRNA labeled with Cy5 at the 3’-end (red sphere). (**B**) Electrophoretic mobility shift assay of FnCas12a-crRNA complexes. WT FnCas12a, FnCas12a helix 1 and BH variants were used in a two-fold excess of protein (20 nM) over crRNA (10 nM, 43 nt). Free RNA migrates into the gel and is separated according to its size, whereas RNA bound to protein stays in the gel pockets. (**C**) Quantification of crRNA-Cas12a complexes from three independent assays. (**D**) Schematic representation of pre-crRNA with an additional nucleotide at the 5’-end (44 nt) labeled with Cy3 at the 5’-end (green sphere). (**E**) Investigation of pre-crRNA processing by FnCas12a h1 variants. WT FnCas12a, FnCas12a helix 1 and BH variants were used in a two-fold excess of protein (20 nM) over crRNA (10 nM). Pre-crRNA processing can be followed by the cleavage of the single labeled 5’ nucleotide. (**F**) Quantification of pre-crRNA-Cas12a cleavage from three independent assays.

FnCas12a processes pre-crRNAs to yield mature crRNAs independently. We, therefore, tested whether crRNA processing is also impaired. To this end, we used a pre-crRNA one nucleotide longer than the mature crRNA (**Figure 3D/E**) with a fluorescent label positioned at the 5’-end. Cleavage of the single nucleotide at the 5’-end of the pre-crRNA can be detected: incubation of pre-crRNA with FnCas12a results in a single fluorescently labeled nucleotide and an unlabeled mature crRNA. We also included an FnCas12a variant mutated in the catalytic site responsible for pre-crRNA processing (mutations H843A K852A K869A, termed HKK mutant in this study). Congruent with the reduced crRNA binding capacity of the Δh1 and ΔBH/Δh1 variants, these FnCas12a variants process the pre-crRNA very inefficiently. For all other FnCas12a only minor or no changes in crRNA binding and pre-crRNA processing were observed.

### Modifications in helix 1 reduce *cis-* and *trans-*cleavage efficiency and alter the cleavage position of FnCas12a

Next, we tested whether helix 1 is important for the DNA nuclease activity of FnCas12a using first linearized and supercoiled plasmid DNA as target DNA (**Figure 4**). Cleavage of linearized plasmid DNA results in two DNA fragments 4400 and 2900 bp in size. Wildtype FnCas12a and the K978A variant cleave linearized plasmid DNA with 100% efficiency under the chosen reaction conditions. The K981P, I960P, and ΔBH variants were less efficient in cleavage and the K981P and ΔBH variants showed a smeared cleavage pattern (**Figure 4A**). The I960P/K981P mutant, as well as the Δh1 and ΔBH/Δh1, did not cleave the plasmid DNA at all (**Figure 4A**). Interestingly, DNA cleavage using supercoiled DNA resulted in DNA cleavage even if the I960P/K981P, Δh1, and ΔBH/Δh1 variants were used (**Figure 4B**). However, these variants were inefficient in cleavage. While nicking of one DNA strand occurred, only a minor fraction of fully cleaved (e.g. cleavage of template and non-template strand) and therefore linearized DNA was detected. However, in the case of the K981P and the BH deletion variants (ΔBH and ΔΔBH), a smear between the supercoiled and linearized cleavage product indicates that these FnCas12a variants cleave the DNA in an imprecise fashion.

**Figure 4.**
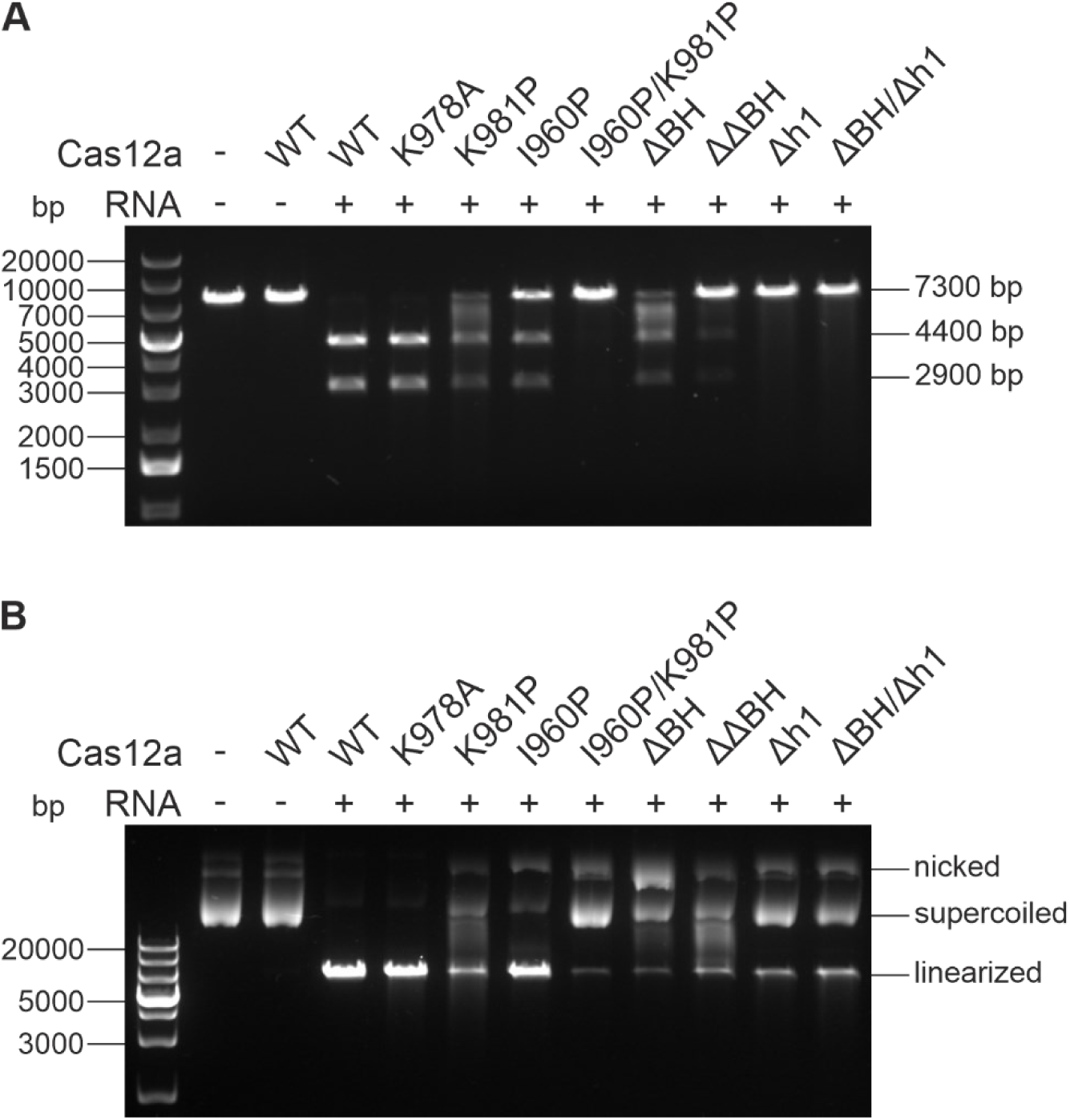
Plasmid cleavage assay of FnCas12a h1 variants. The binary complex (37.5 nM FnCas12a + crRNA) was incubated with 5 nM of target DNA at 37 °C for 1 h. (**A**) The linearized target DNA plasmid (7300 bp) is cleaved into two products (4400 and 2900 bp) and analyzed on a 0.8% agarose 1x TAE gel. (**B**) The supercoiled plasmid DNA is nicked or linearized by FnCas12a variants and analyzed on a 1.5% agarose 1x TAE gel. 1 kb plus DNA ladder.

Using short fluorescently DNA duplexes, we analyzed the cleavage pattern and cleavage accuracy of the FnCas12a variants (**Figure 5**). Cleavage of the DNA by the K978A mutant led to cleavage efficiency as observed for the wildtype with typical cleavage patterns that indicate trimming of the template (TS) and non-template strand (NTS), respectively. As observed in the plasmid cleavage assay, the I960P/K981P mutant was not able to cleave the DNA even though it binds the DNA (**Figure S5**). The single K981P mutant, however, exhibited inefficient cleavage with a cleavage pattern previously observed for the I960P and ΔBH mutants. Disruption of the helical nature of either the BH or helix 1 resulted in a strongly reduced DNA trimming activity especially of the NTS. Even though binding of DNA could not be detected for the Δh1 and ΔBH/Δh1 variant (**Figure S5**), cleavage of DNA could be detected albeit at very low efficiency (**Figure 5**). Notably, the cleavage position at the TS was precise but significantly shifted indicating a cleavage position shifted towards the 3’-end of the TS. In contrast, the cleavage position at the NTS was not altered. We also included mismatched targets in our study. Previously, we showed that FnCas12a variants mutated in the BH are more sensitive to DNA mismatches in the target DNA (24). While the K978A variant performs equally well as the wildtype in cleavage of mismatched DNAs, the K981P, I960P/K981P, Δh1, and ΔBH/Δh1 variants did not cleave any of the mismatched DNAs (**Figure S6**).

**Figure 5.**
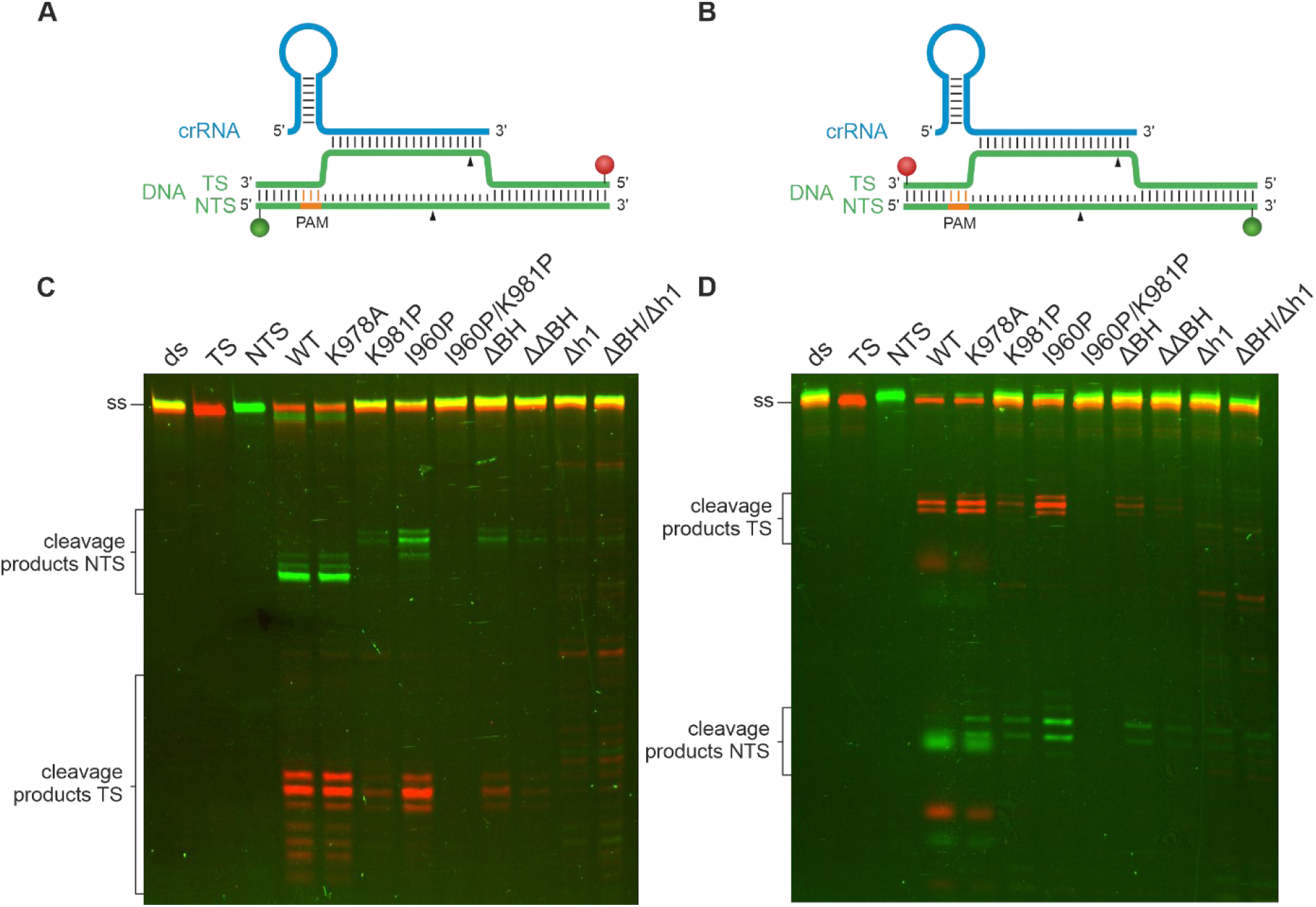
High-resolution cleavage assay of FnCas12a helix 1 variants in comparison to FnCas12a BH variants. The 58 nt double stranded target DNA carries a Cy5 label at the TS and a Cy3 label at the NTS. A 7.5-fold excess of binary complex (Cas12a and crRNA, 750 nM) was used over the target DNA (100 nM). The reaction was incubated for 1 h at 37 °C. (**A**) DNA strands are labeled with Cy3 (green sphere) and Cy5 (red sphere) fluorophores at the 5’-end of the DNA strands, respectively. (**B**) DNA strands are labeled with Cy3 (green sphere) and Cy5 (red sphere) fluorophores at the 3’-end of the DNA strands, respectively. (**C**) Constructs shown in (A) were used. In addition to the previously reported shift of NTS cleavage products of the BH variants the helix 1 deletions variants (Δh1, ΔBH/Δh1) show a shifted cleavage pattern for the TS. (**D**) Constructs shown in (B) were used. The shift in TS cleavage for FnCas12a Δh1 and FnCas12a ΔBH/Δh1 is again observed. FnCas12a I960P/K981P does not show cleavage at all. Samples were analyzed on a denaturing 15% PAA gel.

In addition to *cis*-cleavage of double-stranded DNA, Cas12a proteins are also able to cleave single-stranded DNA in *trans* (26, 29). Therefore, we extended our studies to observe the *trans*-cleavage activity of FnCas12a. To this end, we made use of a fluorescently labeled ssDNA target that also carries a quencher molecule resulting in low fluorescence of the intact DNA strand. *Trans*-cleavage of the ssDNA by Cas12a leads to the separation of fluorophore and quencher and consequently, *trans*-cleavage can be observed over time as an increase in fluorescence (**Figure 6A**). *Trans*-cleavage activity for the wildtype enzyme (k_cat_ = 0.03 s^-1^, K_M_ = 9×10^−7^ M and k_cat_/K_M_ = 3,2×10^4^ M^-1^s^-1^), the K981P variant could be observed with K981P being significantly less catalytically active (k_cat_ = 5.5×10^−5^ s^-1^, K_M_ = 2.2×10^− 7^ M and k_cat_/K_M_ = 2,4×10^3^ M^-1^s^-1^) (**Figure 6B**). All other mutants used in this study did not show any *trans*-cleavage activity. This is also true for FnCas12a variants like W971D or W971F for which we reported efficient *cis*-cleavage activity before (**Figure S7**) (24). We, therefore, conclude that the BH and helix 1 are highly important for *trans*-cleavage.

**Figure 6.**
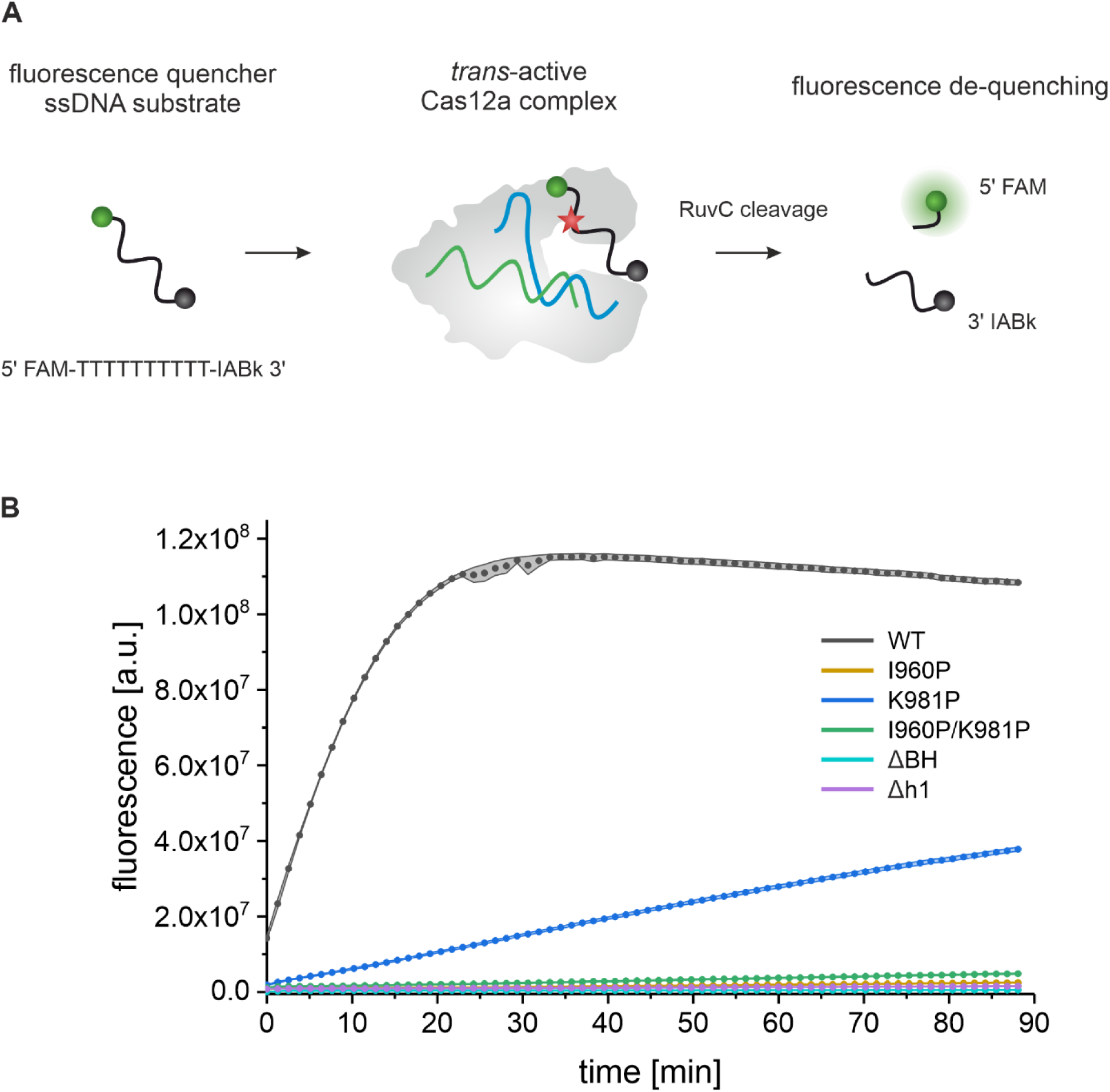
*Trans*-cleavage assay of FnCas12a helix 1 variants in comparison to FnCas12a BH variants. (**A**) The binary complex (100 nM – of premixed 1 µM FnCas12a and 1 µM crRNA) was incubated with an ssDNA activator (5 nM ssDNA) and 100 nM Fluorophore-Quencher (FQ) reporter substrate in a *trans*-cleavage buffer for 20 min. (B) Fluorescence level over time was measured on a plate reader for 60 min. Data points represent three replicates from independent experiments. Error bars indicate the mean ± SEM.

## Discussion

In many respects, the single effector CRISPR-Cas nucleases Cas9 and Cas12a share a comparable structural organization and follow analogous mechanisms throughout their activity cycle (49). Both proteins are divided into two lobes that are connected by a prominent alpha helix called bridge helix. Loading of the crRNA and target DNA is accompanied by closing and re-opening of the two lobes, respectively (11, 15, 50, 23, 33, 34, 40, 45–48). Mutations in the BH of Cas9 influence crRNA and target DNA binding and affect the mismatch tolerance of the enzyme (51). We recently showed that – despite its central position and connecting role – the BH of FnCas12a is not strictly required for the structural integrity of FnCas12a. While the catalytic activity of FnCas12a is reduced when the BH is deleted, it is not fully inactivated. In contrast to Cas9, the BH in Cas12a is connected to a second helix in the RuvC II domain (called helix 1) and structural studies showed that the length of the helices is subject to modulation upon transition from the binary to ternary complex (11, 15, 23, 33, 34, 40). In order to understand whether the rare contact of the helix 1 to the DNA is important to the function of Fncas12a, we tested an FnCas12a variant that is mutated at position K978 – a side chain that contacts the template strand. Mutating the lysine residue to alanine did not result in any loss of activity and hence, the interaction between this amino acid and the DNA is not of importance for the catalytic activity of FnCas12a. We furthermore sought to understand the functional role of helix 1 and made use of BH and helix 1 deletion mutants as well as a double proline mutant (one proline substitution in each helix). Using the BH and helix 1 variants, we surveyed the catalytic activities (RNA processing, *cis-* and *trans*-cleavage of DNA) and conformational transitions of FnCas12a throughout its activity cycle.

Deletion of helix 1 results in an enzyme that cannot cleave linear target DNA, has only minimal activity when using supercoiled plasmid DNA and does not show *trans-*cleavage activity. As shown before, cleavage rates increase when using supercoiled DNA as compared to linear DNA (19, 21, 27, 52, 53). Supercoiled DNA is prone to DNA breathing, which aids R-loop formation. Thus, for FnCas12a variants devoid of the BH or helix 1 that are otherwise cleavage incompetent, rare cleavage events can be observed. However, pre-crRNAs are still processed. Protein stability and single-molecule FRET data indicate that the stability of the helix 1 deletion variant is reduced and that mainly the open conformation of FnCas12a is adopted. Nevertheless, the overall folding of the enzyme appears to be intact. This seems to include the catalytic site in the WED domain that is required for 5’-end cleavage of the pre-crRNA as the pre-crRNA can be processed. It is unlikely that the folding and conformation of the WED domain is affected by helix 1 that is structurally not connected to the active site positioned in the WED domain. Transition into the closed conformation of FnCas12a is not induced upon crRNA binding in the helix 1 deletion variant but the conformational transitions in FnCas12a required for the activation of the catalytic site (11, 12) in the WED domain must be possible. Therefore, the RNA processing activity is not abolished even if helix 1 or the BH are deleted. Similarly, disruption of the helical structure of the BH and/or helix 1 (single proline or double proline mutations in the BH and helix 1) does not significantly impact the pre-crRNA processing step. Especially in the case of the helix 1 deletion variant, we noticed that binding of the crRNA is more strongly impaired than pre-crRNA processing. As the binding site of the mature and pre-crRNA is the same, we speculate that the pre-crRNA might only bind transiently allowing processing but not stable binding of the crRNA. This is in line with the smFRET measurements, in which we do not observe the shift to the high FRET population typically observed for the binary complex. As smFRET measurements are performed at lower nanomolar concentrations, a transient binding event would not lead to a stable complex and hence, we do not observe the binary complex in these measurements. In contrast to pre-crRNA processing, the conformational transitions that are required for loading and cleavage of DNA are prevented if helix 1 is deleted. The role of helix 1 in the stabilization of the overall structure of FnCas12a cannot be compensated by the BH or the crRNA and consequently, the closure of the enzyme upon binding of the crRNA is not possible. In contrast, the deletion of the BH results only in moderate impairments in the structure and function of FnCas12a suggesting that helix 1 is the central helix that couples the Nuc and REC lobe. Notably, the FnCas12a^I960P/K981P^ variant is able to bind DNA and go through all conformational changes that are observed for the wildtype enzyme. However, DNA *cis-* and *trans-*cleavage was not observed. Also, the FnCas12a^K981P^ variant that carries a single proline substitution in helix 1 shows a more drastic reduction in catalytic activity as compared to the variant that carries a single proline substitution in the BH. This underscores the functional importance of helix 1. However, as neither of the helices forms extensive interactions with the crRNA or DNA and because the major conformational transitions in the enzyme are still possible, it has to be assumed that the allosteric activation of the active center in the RuvC domain is prevented in the FnCas12a^I960P/K981P^ variant. Structural studies showed that large and small conformational changes activate the RuvC cleavage site upon R-loop formation (11, 18, 33, 34, 40). Our smFRET data suggest that the large rotational movement of the Rec lobe is not impaired even if the helical nature of the BH and helix 1 is disrupted. Smaller conformational changes also occur in structural elements called the ‘REC linker’ (connecting the REC1 and REC2 domain), the ‘lid’ (part of the RuvC domain), and the ‘finger’ (located in the REC1 domain) (18). In the binary complex, the lid is closed thereby denying access to the active site. Upon ternary complex formation, the lid is found in an open conformation and the catalytic site in the RuvC domain (formed by residues D917, E1006, D1255) is stabilized by polar interactions of lysine 1013 (part of the lid) with aspartate 917. Importantly, the residues that form the lid (residues F1005-K1021 in FnCas12a) are located just four amino acids away from the C-terminal end of helix 1 (helix 1 spans residues K972-N1000). Helix 1 and the BH undergo a tandem movement upon transition from the binary to the ternary complex that results in a shortening of helix 1 and an extension of the BH.

However, replacing the lysine at position 981 by proline or introducing two proline residues in each helix most likely interferes with this tandem movement (**Figure 7**). Most importantly, it seems plausible that impairments in the transition of helix 1 directly impact the closed-to-opening transition of the lid upon ternary complex formation. We suggest that the lid cannot open completely and that the catalytic residues are not accurately aligned for cleavage (one of the catalytic residues is E1006, which is part of the lid) leading to a catalytically inactive enzyme (**Figure 7B**).

**Figure 7.**
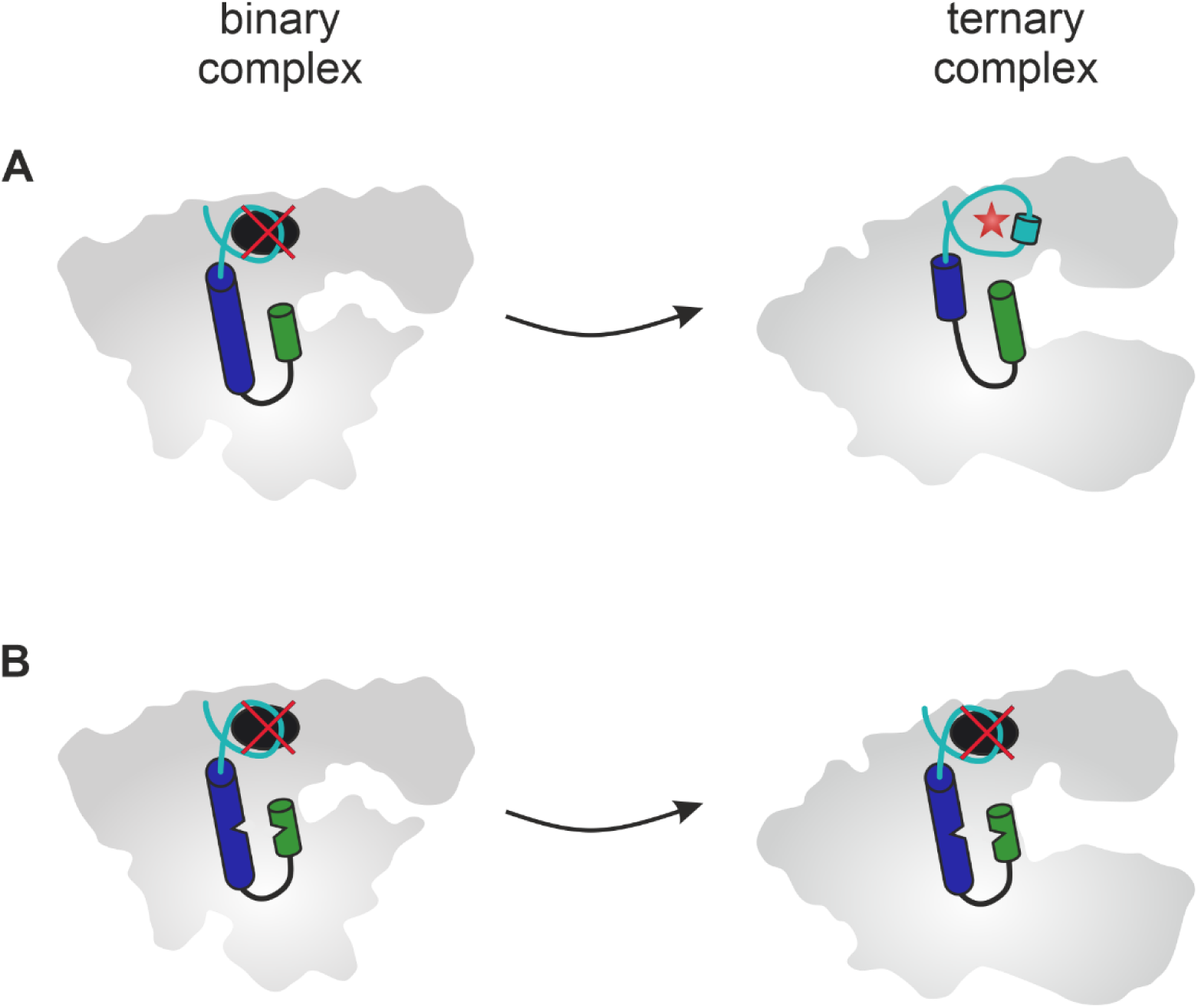
Mechanistic model summarizing the impact of helix 1 on FnCas12a activity. (**A**) Loading of target DNA leads to the formation of the ternary complex. This transition is accompanied by an opening of the enzyme and a tandem movement of helix 1 and the bridge helix (BH), e.g. a shortening of helix 1 (blue cylinder) and extension of the BH (green cylinder). Additionally, the lid domain (cyan) is restructured inducing the formation of a short helix and ultimately the opening and activation of the active site in the RuvC II domain. (**B**) Introduction of proline residues in helix 1 and the BH (kink in the cylinders), respectively, prevents the concerted tandem movement of helix 1 and the BH. The impaired movement of helix 1 is translated to the lid domain that cannot reorder ultimately preventing the opening of the lid. Consequently, the active site is not activated and *cis*- and *trans*-DNA cleavage activity cannot be observed.

Taken together, our study reveals helix 1 (and not the BH) is central to the structure and function of FnCas12a. We furthermore show that the BH, helix 1, and the lid act as a finely tuned structural unit, and distortion of this unit directly impacts the active site of the enzyme via the lid.

## Data availability

The datasets generated during and/or analyzed in this study are available from the corresponding author on reasonable request.

## Acknowledgements

We thank Leonhard Jakob for discussions in the early stages of this project and gratefully thank Andreas Schmidbauer for critical reading of the manuscript and fruitful discussions. We furthermore thank Elisabeth Piechatschek and Elke Papst for technical assistance. The project was undertaken with the assistance of resources and services from the National computational infrastructure (NCI), which is supported by the Australian Government.

## Author contributions

D.G. conceived the study. E.W. purified the proteins and created FnCas12a variants. E.W. performed EMSAs. E.W. and A.N. performed DNA *cis*-cleavage assays. A.N. and G.B. performed and analyzed the DNA *trans*-cleavage assays. G.B. performed the structural modeling. E.W. performed the single-molecule measurements. E.W. analyzed the single-molecule data. E.W. and D.G. wrote the paper. All authors commented on the paper.

