## Supplemental Information for "Allosteric activation of CRISPR-Cas12a requires the concerted movement of the bridge helix and helix 1 of the RuvC II domain"

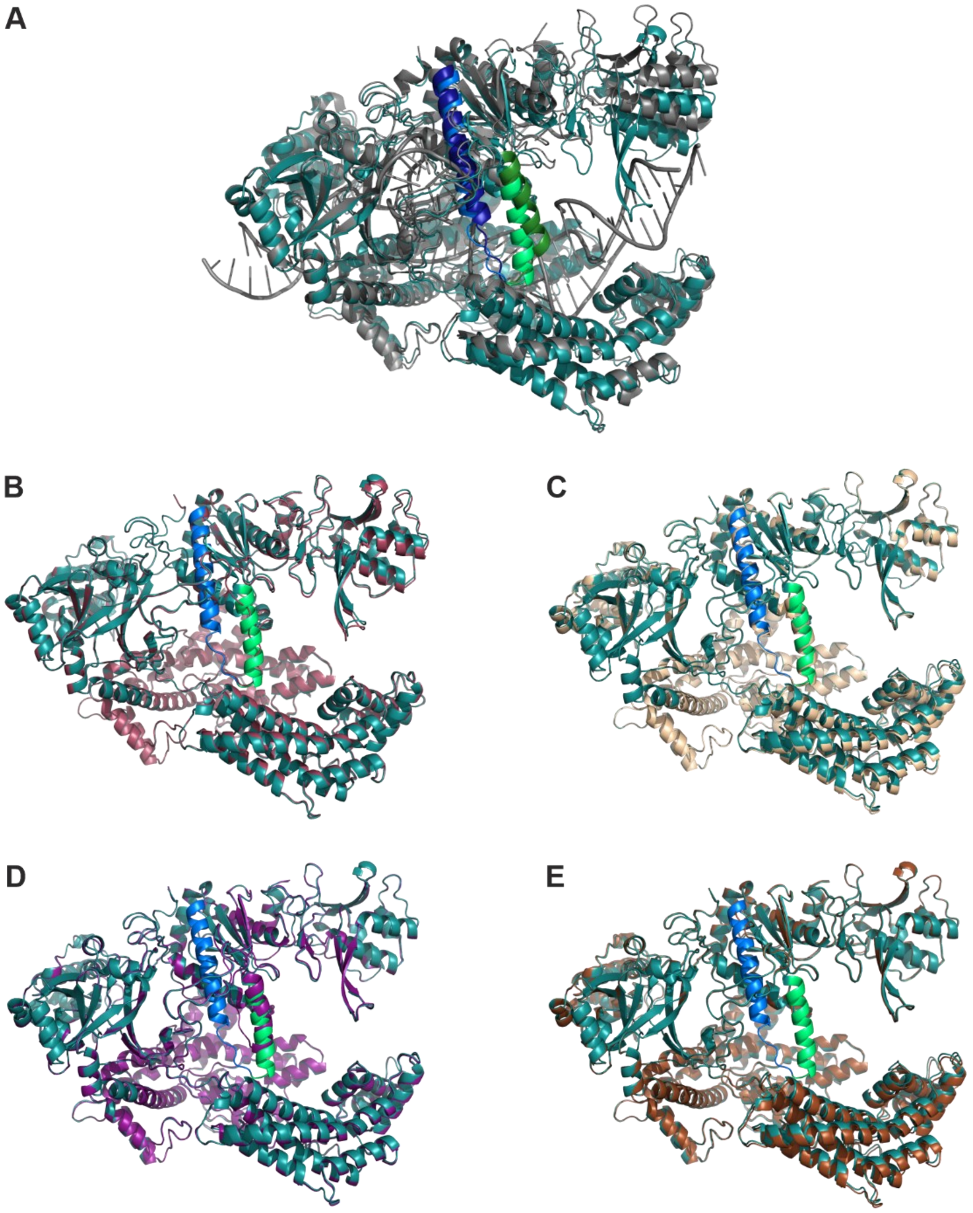


**Figure S1.** Structural alignment of FnCas12a wildtype and FnCas12a variant structures. (**A**) Structural alignment of WT FnCas12a ternary structure (PDB: 6I1K, bound to crRNA and target DNA, grey) and the WT structure predicted by AlphaFold (cyan). The BH is colored in green (6I1K: dark green, AlphaFold: light green), h1 in blue (6I1K: dark blue, AlphaFold: light blue). (**B**) to (**E**) Structural alignment of FnCas12a variants with WT FnCas12a (cyan). FnCas12a structures were predicted by AlphaFold. BH and h1 of the WT structure are colored in light green and light blue, respectively. (**B**) ΔBH (red) (**C**) I960P/K981P (yellow) (**D**) Δh1 (violet) (**E**) ΔBHΔh1 (brown).

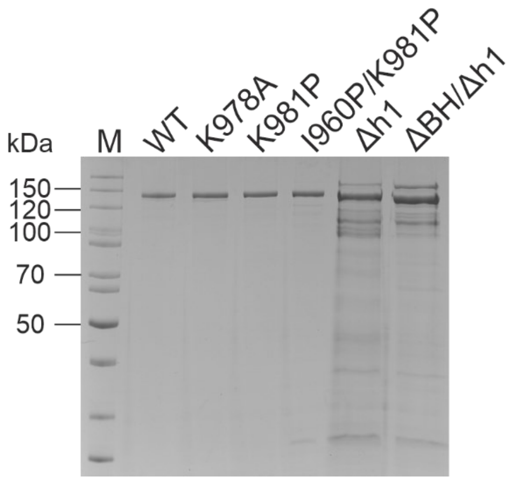


**Figure S2.** SDS-PAGE of purified FnCas12a variants (1.7 µg each, molecular weight of WT FnCas12a: 151.9 kDa). M: PageRuler™ (Thermo Scientific) unstained.

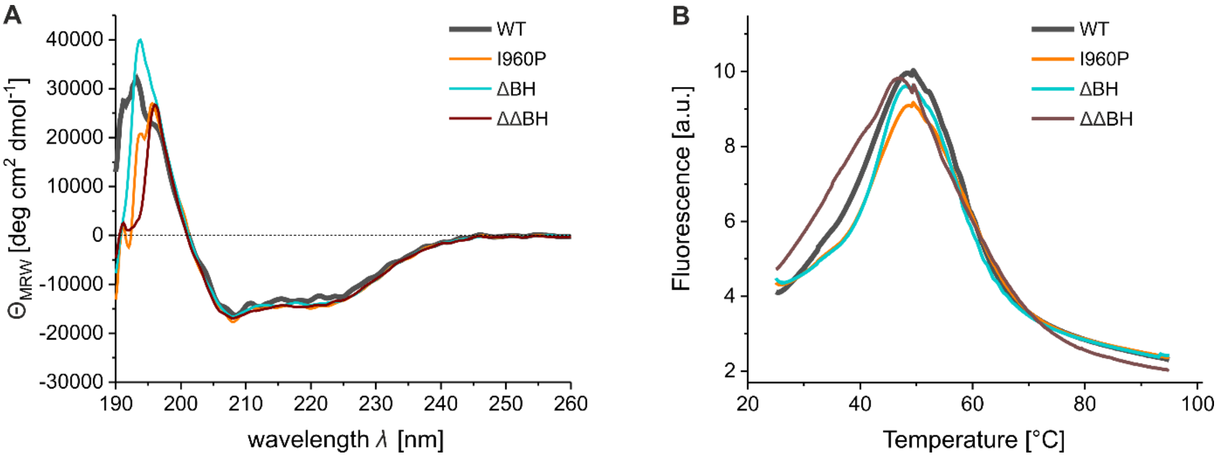


**Figure S3.** CD spectra and melting curves of WT FnCas12a and FnCas12a BH variants. (**A**) Far-UV CD spectra of WT FnCas12a and FnCas12a BH variants (2 µM) at 25 °C. Shown is the average of five replicates. (**B**) Protein Thermal Shift™ (Thermo Scientific) melting curves of WT FnCas12a and FnCas12a BH variants (2 µg) from 25 to 95 °C. Shown is the average of four replicates.

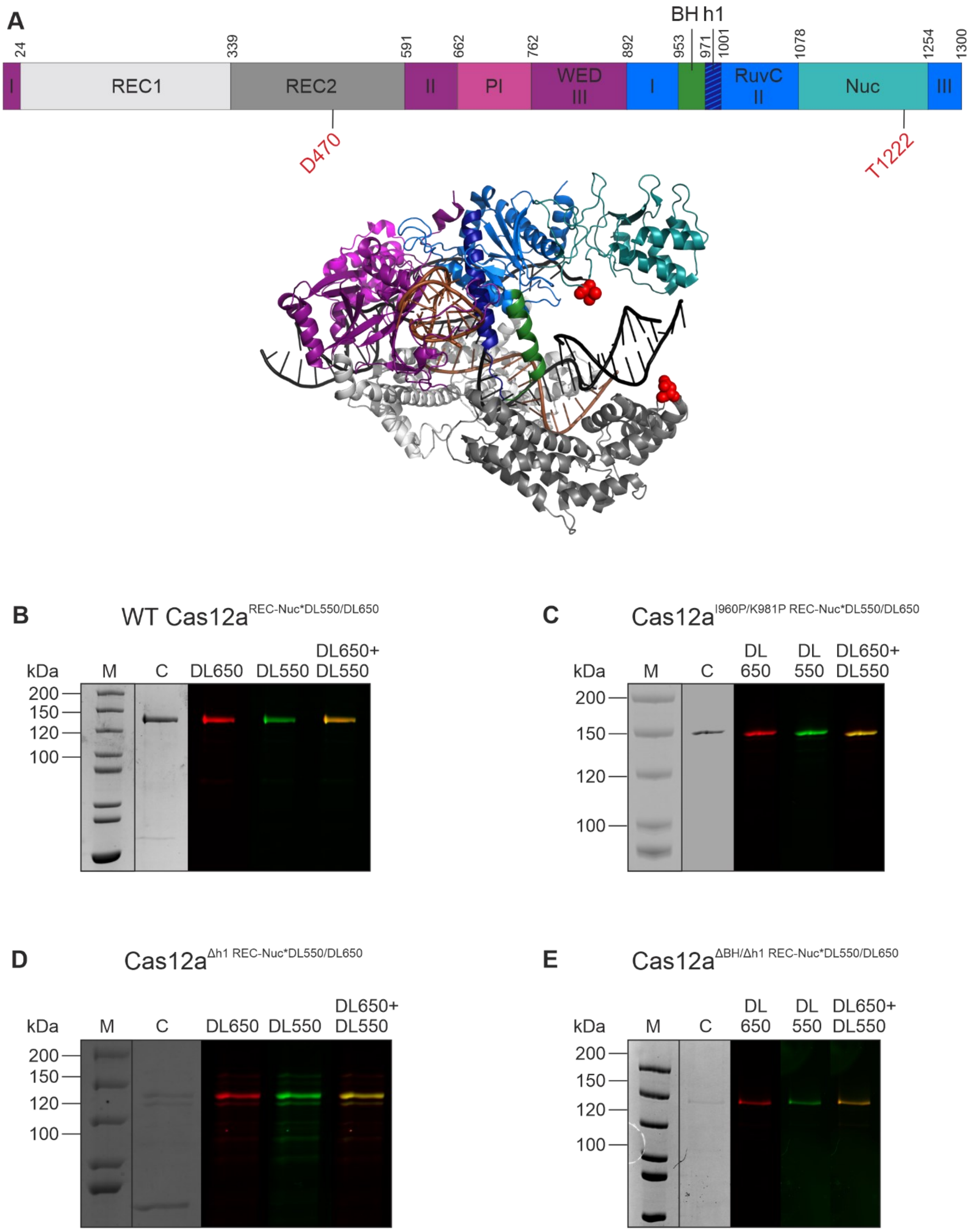


**Figure S4.** (**A**) Domain organization and structure of the FnCas12a ternary complex with labeling sites indicated in red (PDB: 6I1K). (**B**)-(**E**) 10% SDS-PAGE analysis of doubly labeled FnCas12a variants with DyLight 550 (DL550) and DyLight 650 (DL650)) fluorophores. M: PageRuler™ unstained; C: Coomassie stained gel; DL650: fluorescence scan at 625-650 nm excitation for DyLight 650; DL550: fluorescence scan at 520-545 nm excitation for DyLight 550; DL650+DL550: overlay of fluorescence scans of DL650 and DL550.

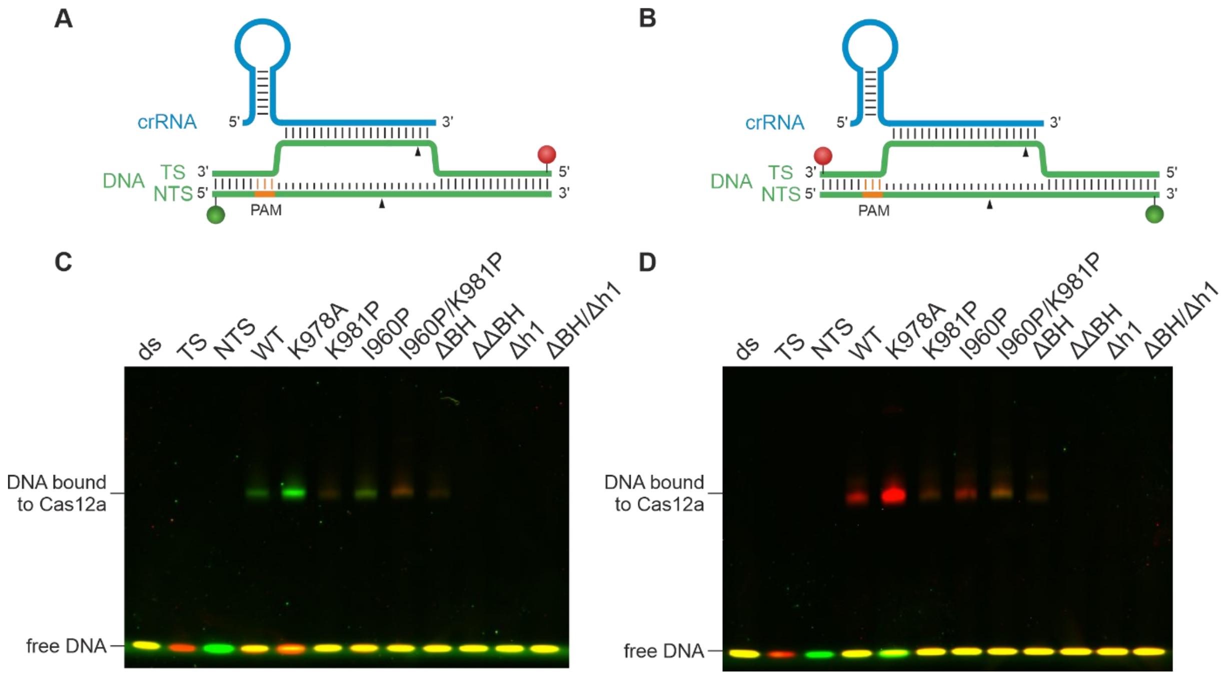


**Figure S5.** Binding behavior of FnCas12a helix 1 variants to target DNA in comparison to FnCas12a BH variants. Electrophoretic mobility shift assays of FnCas12a-crRNA-target DNA complexes. The proteins WT FnCas12a, FnCas12a helix 1 and FnCas12a BH variants and crRNA were used in a 7.5-fold excess (75 nM) over target DNA (10 nM). The DNA (58 nt) is doubly labeled with a Cy3 label (green) at the non-target strand (NTS) and a Cy5 label (red) at the target strand (TS). (**A**) DNA strands are labeled with Cy3 (green sphere) and Cy5 (red sphere) fluorophores at the 5'-ends, respectively. (**B**) DNA strands are labeled with Cy3 (green sphere) and Cy5 (red sphere) fluorophores at the 3'-ends, respectively. (**C**) Constructs shown in (A) were used. Free DNA migrates until the bottom of the gel, protein-bound DNA is shifted and migrates slower. The protein-bound DNA band appears green, as after DNA cleavage, the protein remains bound to the PAM proximal part of the DNA (Cy3). In complexes with less active variants (Cas12a K981P, I960P K981P, ΔBH) the Cy5-labeled DNA is still bound to the protein. (**D**) Constructs shown in (B) were used. Free DNA migrates until the bottom of the gel, protein-bound DNA is shifted and migrates slower. The protein-bound DNA band appears red, as after DNA cleavage, the protein remains bound to the PAM proximal part of the DNA (Cy5). In complexes with less active variants (Cas12a K981P, I960P K981P, ΔBH) the Cy3-labeled DNA is still bound to the protein. FnCas12a variants ΔΔBH, Δh1, and ΔBHh1 show no binding to DNA.

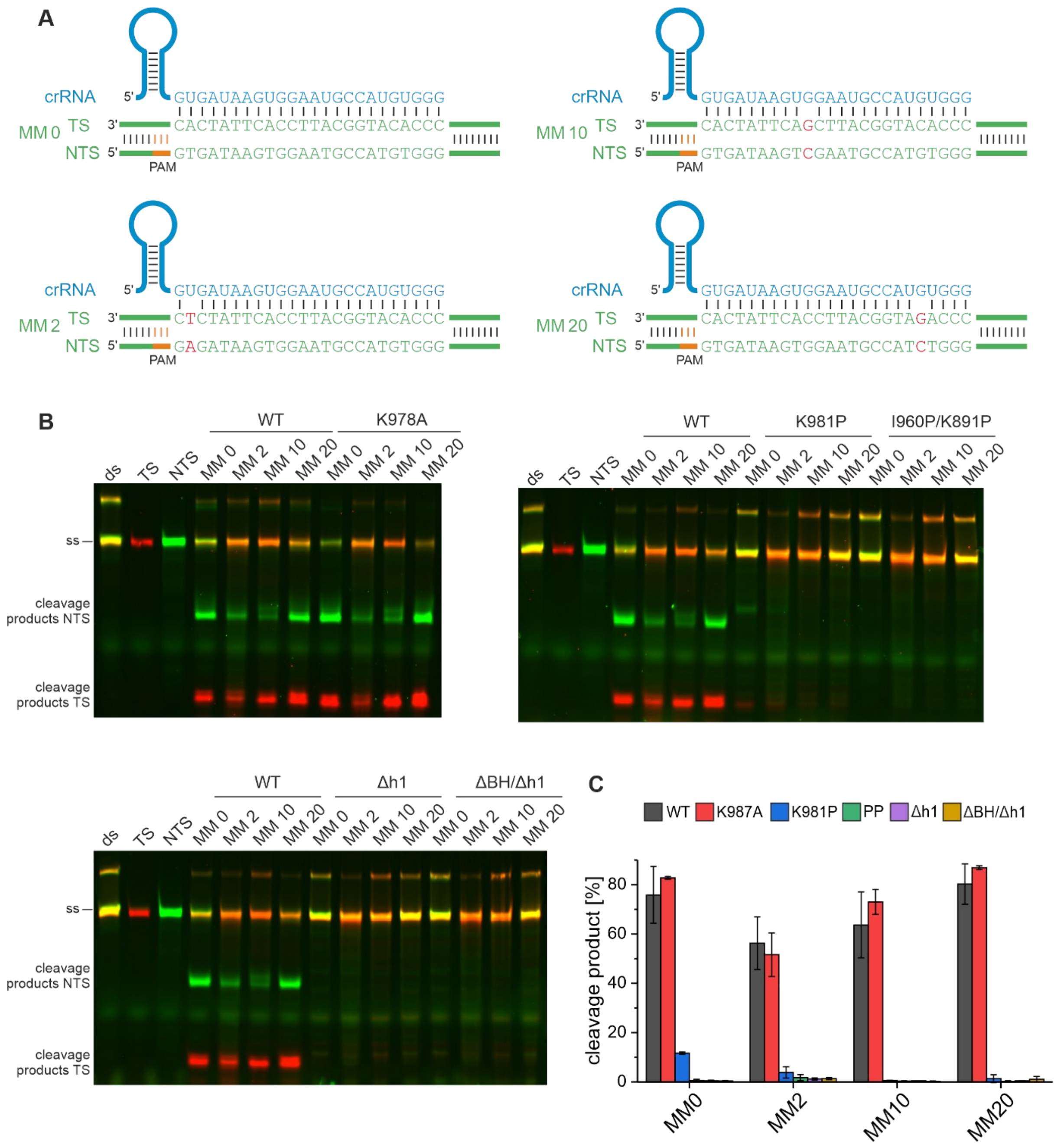


**Figure S6.** DNA cleavage specificity of FnCas12a helix 1 variants at mismatched target DNA. (**A**) Representation of the analyzed target DNA substrates with full complementarity between crRNA and target DNA (MM 0), and mismatches at position 2 (MM 2), position 10 (MM 10), or position 20 (MM 20) between crRNA and target DNA. The DNA constructs (58 nt) are 5'-labeled with Cy3 at the NTS and Cy5 at the TS. (**B**) Cleavage assays of FnCas12a helix 1 variants and crRNA in 7.5-fold excess (75 nM) to mismatched target DNA (10 nM). The reaction was performed for 1 h at 37 °C and analyzed with denaturing 15% PAA gels. (**C**) Quantification of the NTS cleavage efficiency of FnCas12a helix 1 variants. FnCas12a K978A shows comparable cleavage activity to WT FnCas12a, FnCas12a K981P is only marginally active at the fully complementary and the MM 2 target construct. The variants FnCas12a I960P K981P, Δh1, and ΔBHh1 show no cleavage activity.

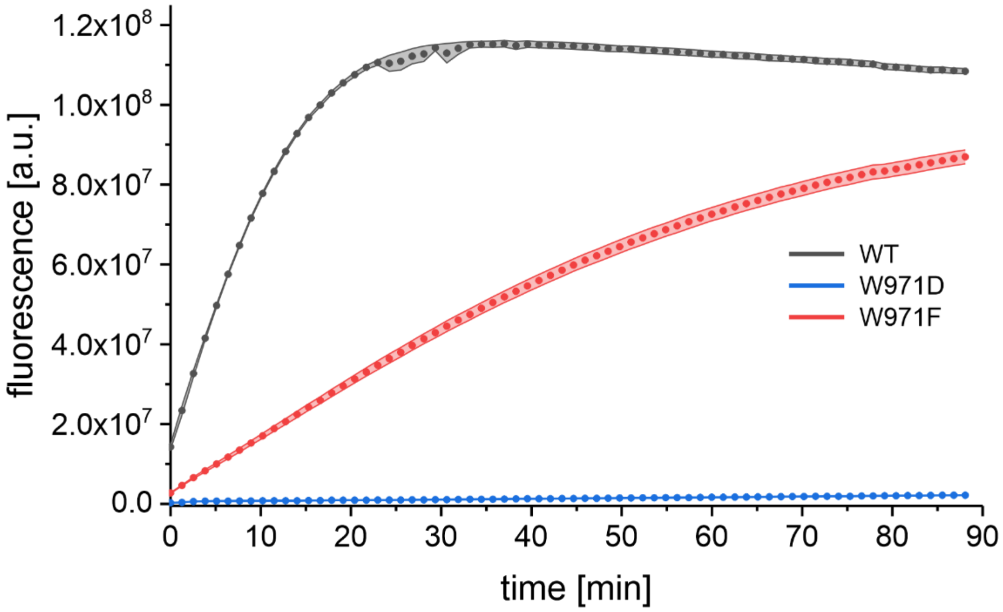


**Figure S7.** *Trans*-cleavage activity of FnCas12a BH variants W971D and W971F in comparison to the FnCas12a WT enzyme.

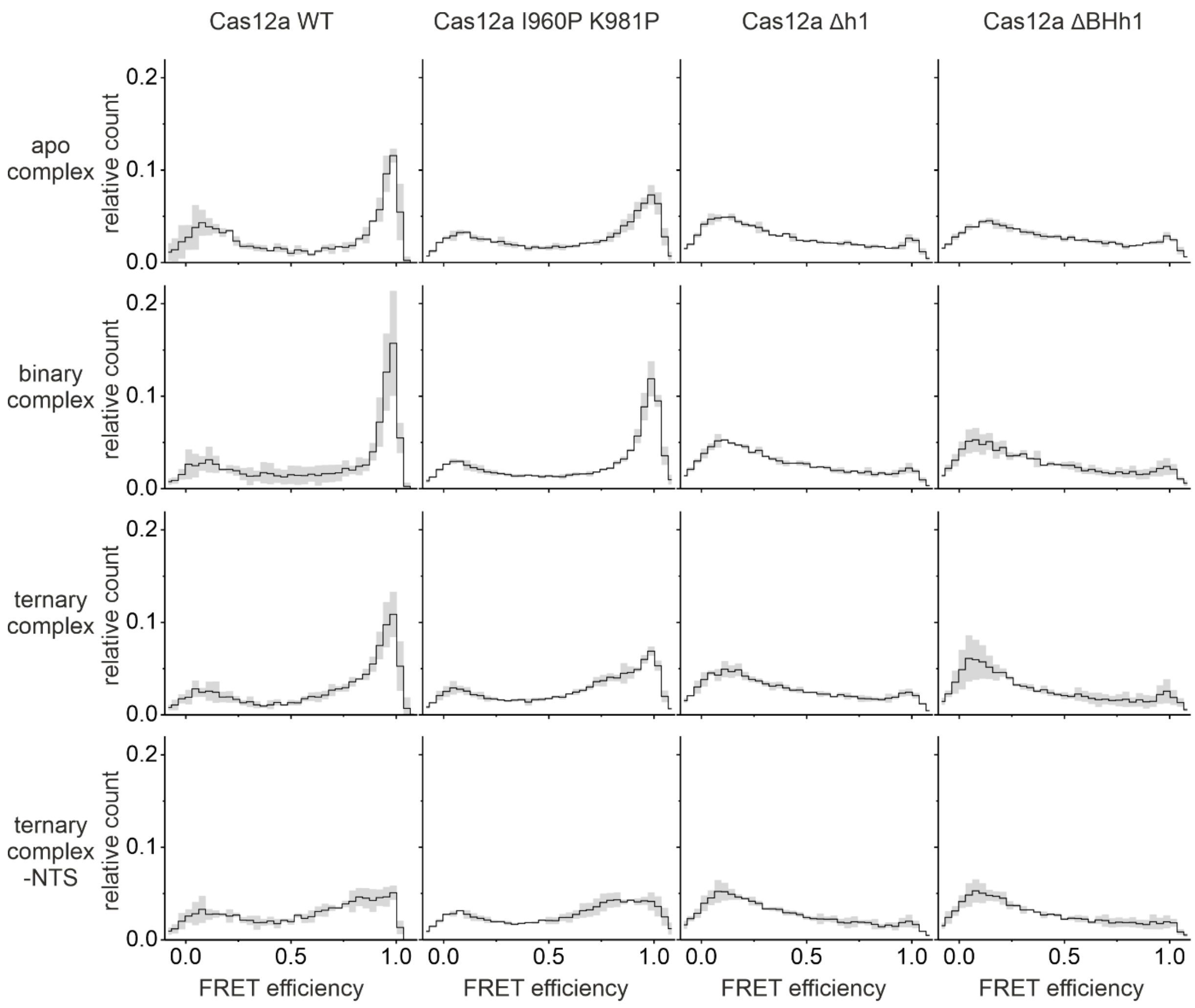


**Figure S8.** FRET efficiency histograms with mean FRET efficiency (black) and standard deviation (grey) of three measurements are shown for WT FnCas12a$^{REC\text{-}Nuc*DL550/DL650}$ and the FnCas12a$^{REC\text{-}Nuc*DL550/DL650}$ helix 1 variants. Histograms represent the apo enzyme, the binary complex (1 nM crRNA), the ternary complex (1 nM crRNA, 1 nM target DNA), and the ternary complex without NTS (1 nM crRNA, 1 nM TS DNA).

**Table S1.** Sequences of mutagenesis primer.

| | | Sequence 5' → 3' |
|---|---|---|
| **I960P** | fw | CATGATAAGCTTGCTGCACCAGAGAAAGATAGGGATTC |
| | rev | GAATCCCTATCTTTCTCTGGTGCAGCAAGCTTATCATG |
| **K978A** | fw | GGAAAAAGATAAATAACATCGCTGAGATGAAAGAGGG |
| | rev | CCCTCTTTCATCTCAGCGATGTTATTTATCTTTTTCC |
| **K981P** | fw | GATAAATAACATCAAAGAGATGCCTGAGGGCTATCTATCTCAGG |
| | rev | CCTGAGATAGATAGCCCTCAGGCATCTCTTTGATGTTATTTATC |
| **ΔBH** | fw | GGTAATGATAGAATGAAAACAAACGACTGGAAAAAGATAAATAAC |
| (Y953-K969) | rev | GTTATTTATCTTTTTCCAGTCGTTTGTTTTCATTCTATCATTACC |
| **ΔΔBH** | fw | GGTAATGATAGAATGAAAACAAACAAAAAGATAAATAACATCAAAGAG |
| (Y953-W971) | rev | CTCTTTGATGTTATTTATCTTTTTGTTTGTTTTCATTCTATCATTACC |
| **Δh1** | fw | GGGATTCAGCTAGGAAAGACTGGGCTATTGTGGTTTTTGAGGATTTAAATTTTGG |
| (K972-N1000) | rev | CCAAAATTTAAATCCTCAAAAACCACAATAGCCCAGTCTTTCCTAGCTGAATCCC |
| **ΔBHh1** | fw | GGTAATGATAGAATGAAAACAAACGCTATTGTGGTTTTTGAGGATTTAAATTTTGG |
| (Y953-N1000) | rev | CCAAAATTTAAATCCTCAAAAACCACAATAGCGTTTGTTTTCATTCTATCATTACC |
| **D470Stop** | fw | GCATAGAGATATATAGAAACAGTGTAGG |
| | rev | CCTACACTGTTTCTATATATCTCTATGC |
| **T1222Stop** | fw | CAAATGCGTAACTCAAAATAGGGTACTGAGTTAGATTATC |
| | rev | GATAATCTAACTCAGTACCCTATTTTGAGTTACGCATTTG |

**Table S2.** Sequences of RNA and DNA oligos.

| | | Sequence 5' → 3' |
|---|---|---|
| **target DNA (MM 0)** | TS | ACTCAATTTTGACAGCCCACATGGCATTCCACTTATCACTAAAGGCAT (1)<br>CCTTCCACGT |
| | NTS | ACGTGGAAGGATGCCTTTAGTGATAAGTGGAATGCCATGTGGGCTGTC (1)<br>AAAATTGAGT |
| **target DNA MM 2** | TS | ACTCAATTTTGACAGCCCACATGGCATTCCACTTATCTCTAAAGGCAT<br>CCTTCCACGT |
| | NTS | ACGTGGAAGGATGCCTTTAGAGATAAGTGGAATGCCATGTGGGCTGTC<br>AAAATTGAGT |
| **target DNA MM 10** | TS | ACTCAATTTTGACAGCCCACATGGCATTCGACTTATCACTAAAGGCAT<br>CCTTCCACGT |
| | NTS | ACGTGGAAGGATGCCTTTAGTGATAAGTCGAATGCCATGTGGGCTGTC<br>AAAATTGAGT |
| **target DNA MM 20** | TS | ACTCAATTTTGACAGCCCAGATGGCATTCCACTTATCACTAAAGGCAT<br>CCTTCCACGT |
| | NTS | ACGTGGAAGGATGCCTTTAGTGATAAGTGGAATGCCATCTGGGCTGTC<br>AAAATTGAGT |
| **T7 DNA for crRNA** | T7 | CCCACATGGCATTCCACTTATCACATCTACAACAGTAGAAATTCCCTA<br>TAGTGAGTCGTATTATCGATC |
| **T7 promotor oligo** | T7 | GATCGATAATACGACTCACTATAGGG |
| **crRNA** | | AAUUUCUACUGUUGUAGAUGUGAUAAGUGGAAUGCCAUGUGGG |
| **pre-crRNA** | | UAAUUUCUACUGUUGUAGAUGUGAUAAGUGGAAUGCCAUGUGGG |
| **crRNA *trans*-cleavage** | | UAAUUUCUACUGUUGUAGAUCUGAUGGUCCAUGUCUGUUACUCG |
| **ssDNA activator** | | CCCGGTGTCACGCCACTTGACAGGCGAGTAACAGACATGGACCATCAG<br>GAAACATTAACGTACTGATGTTAACAGCTGACCCAATAAGTGGCAGAG |
| **FQ** | | FAM-TTTTTTTTTT-IABk |

**Table S3.** Distances between D470 and T1222 in variant structures predicted by AlphaFold2.

| | |
|---|---|
| WT | 31.5 Å |
| K981P | 31.8 Å |
| I960P | 32.3 Å |
| I960P/K981P | 32.6 Å |
| ΔBH | 31.2 Å |
| ΔΔBH | 33.8 Å |
| Δh1 | 32.1 Å |
| ΔBH/Δh1 | 32.5 Å |

1. Swarts,D. and Jinek,M. (2018) Mechanistic insights into the *cis*- and *trans*-acting deoxyribonuclease activities of Cas12a. *Mol. Cell*, **73**, 589-600.e4.
